# Differential Response of Digesta- and Mucosa-Associated Intestinal Microbiota to Dietary Black Soldier Fly (*Hermetia illucens*) Larvae Meal in Seawater Phase Atlantic Salmon (*Salmo salar*)

**DOI:** 10.1101/2020.05.08.083899

**Authors:** Yanxian Li, Leonardo Bruni, Alexander Jaramillo-Torres, Karina Gajardo, Trond M. Kortner, Åshild Krogdahl

## Abstract

Intestinal digesta is commonly used for studying responses of microbiota to dietary shifts, yet evidence is accumulating that it represents an incomplete view of the intestinal microbiota. In a 16-week seawater feeding trial, Atlantic salmon (*Salmo salar*) were fed either a commercially-relevant reference diet or an insect meal diet containing 15% black soldier fly (*Hermetia illucens*) larvae meal. The digesta- and mucosa-associated distal intestinal microbiota were profiled by 16S rRNA gene sequencing. Regardless of diet, we observed substantial differences between digesta- and mucosa-associated intestinal microbiota. Microbial richness and diversity were much higher in the digesta than the mucosa. The insect meal diet altered the distal intestinal microbiota resulting in higher microbial richness and diversity. The diet effect, however, depended on the sample origin. Digesta-associated intestinal microbiota showed more pronounced changes than the mucosa-associated microbiota. Lastly, multivariate association analyses identified two mucosa-enriched taxa, *Brevinema andersonii* and unclassified *Spirochaetaceae*, associated with the expression of genes related to immune responses and barrier function in the distal intestine, respectively. Overall, our data clearly indicate that responses in digesta- and mucosa-associated microbiota to dietary inclusion of insect meal differ, with the latter being more resilient to dietary changes.

## Introduction

The reduction in the sequencing costs and the advent of bioinformatics in the past decade have enabled in-depth taxonomic and functional profiling of microbial communities from diverse environments at an unprecedented scale. Recent advances in the microbiome studies have shed light on the role of intestinal microbiota in a wide spectrum of host physiological processes and development of diseases, such as helping digestion and absorption (1), modulating lipid metabolism and energy balance (2, 3), dialoguing with the central nervous system through the so-called microbiota-gut-brain axis (4) and being a risk factor or a therapeutic intervention of inflammatory bowel disease (5–8). Diet is one of the key factors in shaping the intestinal microbiota. While long-term dietary habits have a considerable effect on the structure and activity of host intestinal microbiota (9–11), short-term dietary change also alters the intestinal microbiota in a rapid and reproducible way (12). Different dietary components selectively promote or suppress the growth of certain microbial clades, which in turn could inflict important effects on the host health and disease resistance (13, 14).

Atlantic salmon (*Salmo salar*) is the most produced seawater fish species and one of the most economically important farmed fish worldwide (15). While Atlantic salmon are strictly carnivorous in the wild, farmed Atlantic salmon have experienced a substantial shift in the diet composition due to a limited supply of marine ingredients. Marine ingredients used for Norwegian farmed Atlantic salmon have gradually been replaced by plant sources, decreasing from 90% in 1990 to 25% in 2016 (16). Due to concerns on the economic, environmental and social sustainability of the current raw materials for Atlantic salmon farming (15), more sustainable alternative feed ingredients, such as insects (17) and yeasts (18), have been developed and used. The use of alternative feed ingredients may not only affect the nutrient utilization, fish growth, health, welfare and product quality, but also intestinal microbiota in Atlantic salmon (19–21). While studies in mammals and fish have revealed substantial differences between the digesta- and mucosa-associated intestinal microbiota (19, 22–25), most studies investigating diet effects on the intestinal microbiota of fish have sampled the digesta only or a mixture of digesta and mucosa. Evidence is accumulating that digesta- and mucosa-associated intestinal microbiota in fish respond differently to dietary changes (19, 26–29). Profiling only one of or a mixture of digesta- and mucosa-associated microbiota may obscure the response of intestinal microbiota to dietary changes.

Characterizing intestinal microbiota and its associations with host responses is an essential step towards identifying key microbial clades promoting fish health and welfare. Ultimately, a milestone in the fish microbiota research would be knowing how to selectively manipulate the microbiota to improve the growth performance, disease resistance and health status of farmed fish. The main aims of the work presented herein were (i) to compare distal intestinal microbiota of Atlantic salmon fed a commercially relevant diet and an insect meal-based diet, (ii) to further explore the dissimilarity between digesta- and mucosa-associated microbiota and the differences in their response to dietary changes, and (iii) to identify associations between microbial clades and host responses. This work was part of a larger study consisting of a freshwater and seawater feeding trial that aimed to investigate the nutritional value and possible health effects for Atlantic salmon of a protein-rich insect meal produced from black soldier fly (*Hermetia illucens*) larvae. The present work focuses on the intestinal microbiota in seawater phase Atlantic salmon fed an insect meal diet containing 15% black soldier fly larvae meal for 16 weeks. Results on feed utilization, growth performance, fillet quality, intestinal histomorphology and gene expression have been reported elsewhere (30–32).

## Results

Hereafter, different sample groups are named based on the combination of diet (REF vs. IM) and sample origin (DID vs. DIM). Hence, in addition to the extraction blanks, library blanks and mock, we have four different sample types, i.e., REF-DID, REF-DIM, IM-DID and IM-DIM.

### qPCR

Since Cq values of most mucosa DNA templates were out of the linear range of the standard curve, the raw Cq value was used as a proxy of 16S rRNA gene quantity in the diluted DNA templates (Fig. S1). On average, REF-DID showed the highest 16S rRNA gene quantities (mean Cq = 24.7), followed by the mock (mean Cq = 26.1) and IM-DID (mean Cq = 28.4). Irrespective of diet, mucosa DNA templates (REFDIM, IM-DIM) showed similar 16S rRNA gene quantities (mean Cq = 30) that were close to extraction blanks (mean Cq = 32.4).

### Taxonomic composition

All the eight bacterial species included in the mock were successfully identified at genus level with *Enterococcus faecalis*, *Lactobacillus fermentum*, *Listeria monocytogenes* and *Staphylococcus aureus* further being annotated at the species level (Fig. S2A). At the genus level, the average Pearson’s r between the expected and observed taxonomic profile of the mock was 0.33, whereas the Pearson’s r between the observed taxonomic profile of the mock was 0.98. The relative abundance of most Grampositive bacteria, *Listeria monocytogenes*and *Enterococcus faecalis* in particular, were underestimated. In contrast, the relative abundance of Gram-negative bacteria was overestimated. Most ASVs (97.5% - 99.9%) in the extraction and library blanks were classified as *Pseudomonas* (Fig. S2B), which was the main contaminating taxon removed from the biological samples along with other contaminants including *Curtobacterium*, *Jeotgalicoccus*, *Modestobacter*, *Cutibacterium*, *Hymenobacter*, *Brevundimonas*, *Micrococcus*, *Sphingomonas*, *Devosia*, *Sphingomonas aurantiaca* and *Marinobacter adhaerens*. The exact sequence of the contaminating ASVs and their relative abundance in the extraction and library blanks are available in Table S1.

The taxonomic composition of mucosa samples showed higher similarity than that of the digesta samples, which were more diet-dependent (Fig. 1). At the phylum level, the dominant taxa of mucosa samples for both diets were *Spirochaetes* (REF-DIM, 72 ± 34.6 %; IM-DIM, 47 ± 35.2 %) (mean ± S.D.), *Proteobacteria* (REF-DIM, 21 ± 34.1 %; IM-DIM, 23 ± 34.1 %), *Firmicutes* (REF-DIM, 1 ± 2.8 %; IM-DIM, 11 ± 13.5 %), *Tenericutes* (REF-DIM, 4 ± 8 %; IM-DIM, 8 ± 18.8 %) and *Actinobacteria* (REF-DIM, 1 ± 3.4 %; IM-DIM, 9 ± 8.7 %). For digesta samples, the dominant taxa of REF-DID were *Tenericutes* (33 ± 23.1 %), *Proteobacteria* (31 ± 29.9 %), *Firmicutes* (25 ± 21.1 %) and *Spirochaetes* (9 ± 12.9 %), whereas IM-DID was dominated by *Firmicutes* (45 ± 16.9 %), *Actinobacteria* (25 ± 9.5 %), *Proteobacteria* (17 ± 27.8 %), *Tenericutes* (7 ± 8.8 %) and *RsaHF231* (4 ± 1.5 %) (Fig. 1A). At the genus level, the dominant taxa of mucosa samples for both diets were *Brevinema* (REF-DIM, 52 ± 40.1 %; IMDIM, 25 ± 35 %), unclassified *Spirochaetaceae* (REF-DIM, 20 ± 31.8 %; IM-DIM, 22 ± 31.4 %), *Aliivibrio* (REF-DIM, 18 ± 33.5 %; IM-DIM, 18 ± 35.3 %) and *Mycoplasma* (REF-DIM, 4 ± 8 %; IM-DIM, 8 ± 18.8 %). For digesta samples, the dominant taxa of REF-DID were *Mycoplasma* (33 ± 23.1 %), *Aliivibrio* (20 ± 32.3 %), *Photobacterium* (10 ± 12.6 %), *Brevinema* (6 ± 12.5 %) and *Lactobacillus* (5 ± 4 %), whereas IM-DID was dominated by *Aliivibrio* (15 ± 28.2 %), unclassified *Lactobacillales* (14 ± 6 %), *Corynebacterium 1* (13 ± 5 %), *Bacillus* (8 ± 3.4 %), *Mycoplasma* (7 ± 8.8 %) and *Actinomyces* (5 ± 2 %) (Fig. 1B).

**Fig. 1.**
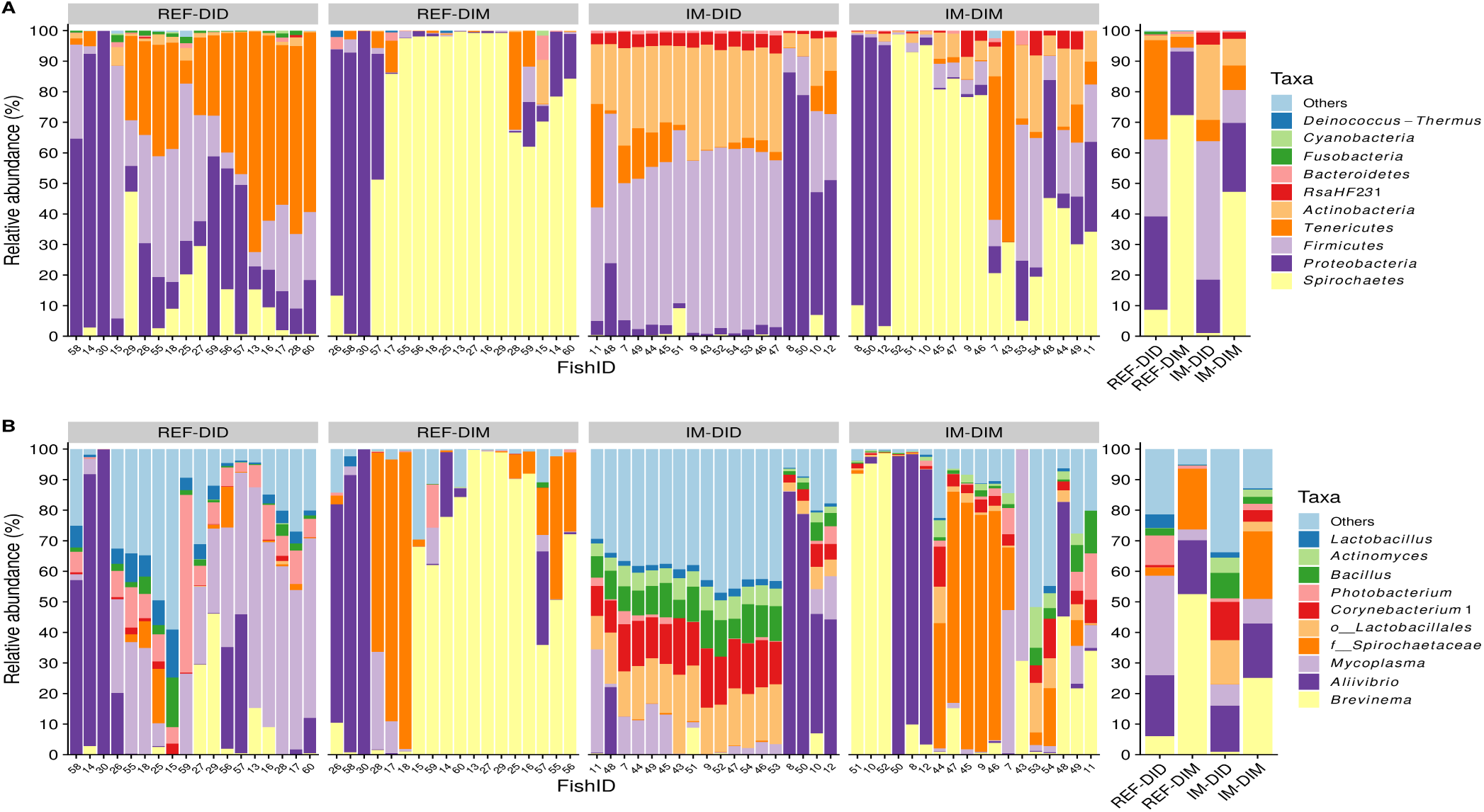
Top 10 most abundant taxa of all samples at phylum (A) and genus (B) level. The samples are grouped by sample origins and dietary treatments. The mean relative abundance of each taxon within the same sample type is displayed on the right side. *o*_, order; *f*_, family; REF, reference diet; IM, insect meal diet; DID, distal intestine digesta; DIM, distal intestine mucosa.

### Core microbiota

In total, 108 taxa were identified as core microbiota based on their prevalence in each sample type (Fig.2; Table S2). Specifically, *Aliivibrio*, *Brevinema andersonii*, and *Mycoplasma* were identified as core microbiota for all the sample types, the latter two being universally present in all the samples. Additionally, ten taxa were identified as core microbiota for digesta samples (REF-DID and IM-DID), which included *Bacillus*, *Corynebacterium 1*, *Lactobacillus* (*L. aviaries*, *L. fermentum* and two unclassified species), *Leuconostoc*, *Parageobacillus toebii*, *Ureibacillus* and *Weissella*. No additional core microbiota taxa were identified for the mucosa samples (REF-DIM and IM-DIM). *Actinomyces*, *Corynebacterium 1*, *Corynebacterium aurimucosum* ATCC 70097, *Microbacterium* and unclassified *RsaHF23* were the additional core microbiota taxa identified for fish fed the insect meal diet (IM-DID and IM-DIM), whereas no additional core microbiota taxa were identified for fish fed the reference diet (REF-DID and REF-DIM). Lastly, 86 taxa were found to be more prevalent in IM-DID than in any other sample type.

**Fig. 2.**
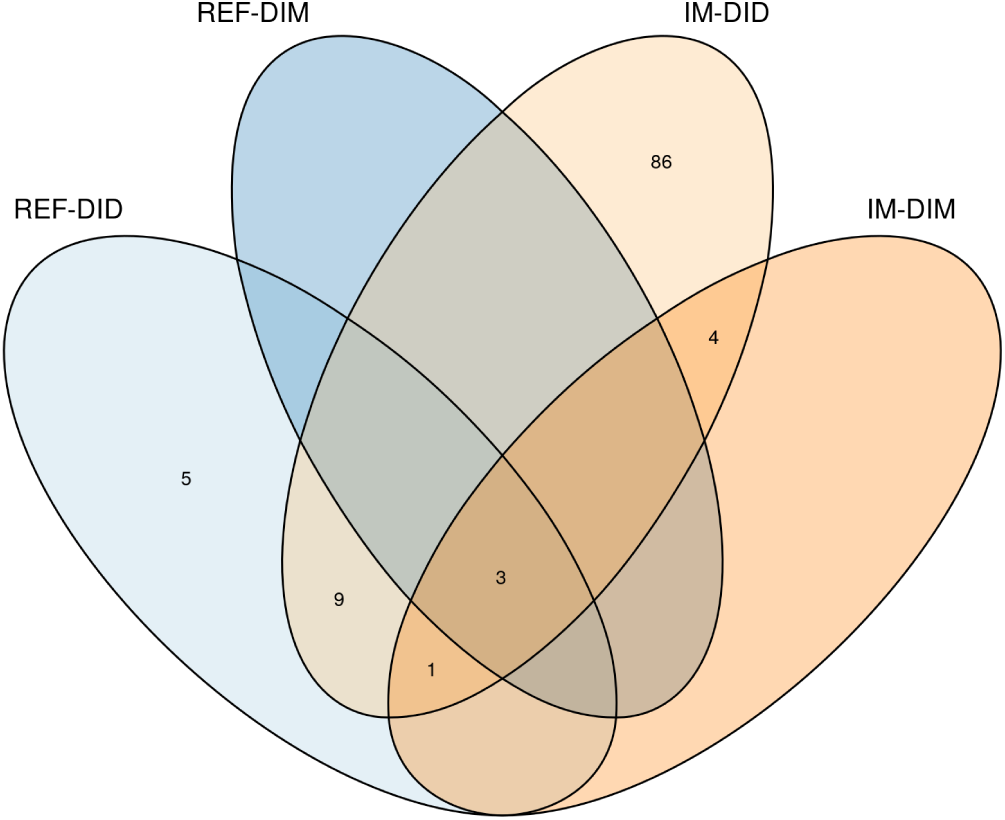
Venn’s diagram showing the shared and unique core microbiota in each sample type. The core microbiota was computed using a prevalence threshold of 80%. REF, reference diet; IM, insect meal diet; DID, distal intestine digesta; DIM, distal intestine mucosa.

### Alpha-diversity

Regardless of diet, all the alpha-diversity indices were higher in digesta samples than mucosa samples (*p* < 0.05) (Fig. 3). Independent of sample origin, all the alpha-diversity indices were higher in fish fed the IM diet than those fed the REF diet (*p* < 0.05). A significant interaction between the diet and sample origin effect was detected for the observed species (*p* = 0.031) and Faith’s phylogenetic diversity (*p* = 0.002), both of which showed a stronger diet effect in digesta samples than mucosa samples.

**Fig. 3.**
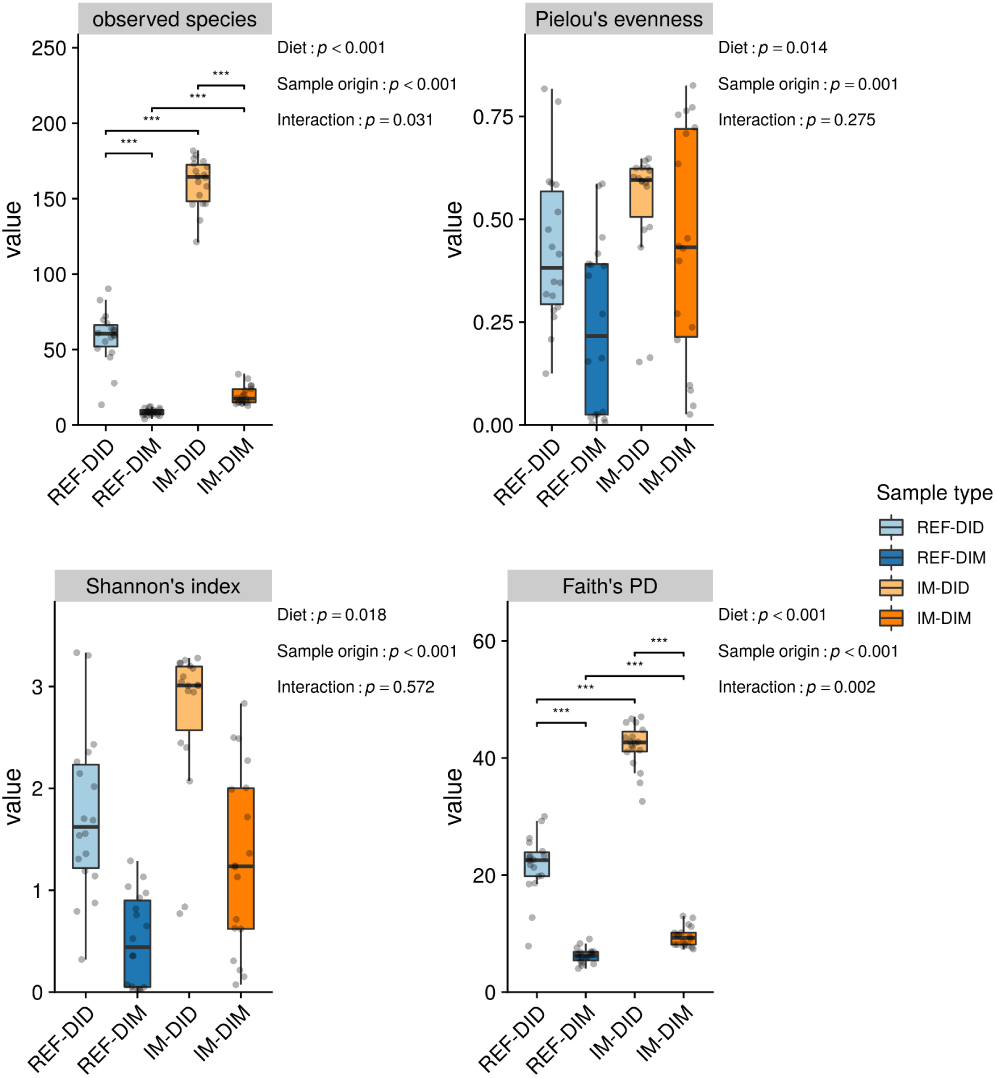
The sample origin and diet effects on the alpha-diversity of distal intestinal microbiota in seawater phase Atlantic salmon. The *p* value of the main effects and their interaction are displayed on the top-right corner of each sub-plot. Asterisks denote statistically significant differences (*, *p* < 0.05; **, *p* < 0.01; ***, *p* < 0.001). PD, phylogenetic diversity; REF, reference diet; IM, insect meal diet; DID, distal intestine digesta; DIM, distal intestine mucosa.

### Beta-diversity

The PCoA plots built on the Jaccard and unweighted UniFrac distance matrix showed clear separations of samples belonging to different dietary groups and sample origins (Fig. 4A-B). However, the average distance between samples from different dietary groups was dependent on sample origin. Specifically, mucosa samples from different dietary groups formed clusters close to each other, whereas digesta samples from different dietary groups were far apart. The PCoA plots built on the Aitchison and PHILR transformed Euclidean distance matrix also showed separations of samples belonging to different dietary groups and sample origins (Fig. 4C-D). Again, the average distance between samples from different dietary groups was dependent on sample origin. Mucosa samples from different dietary groups formed clusters boarding (Fig. 4C) or overlapping (Fig. 4D) each other, whereas digesta samples from different dietary groups were more clearly separated.

**Fig. 4.**
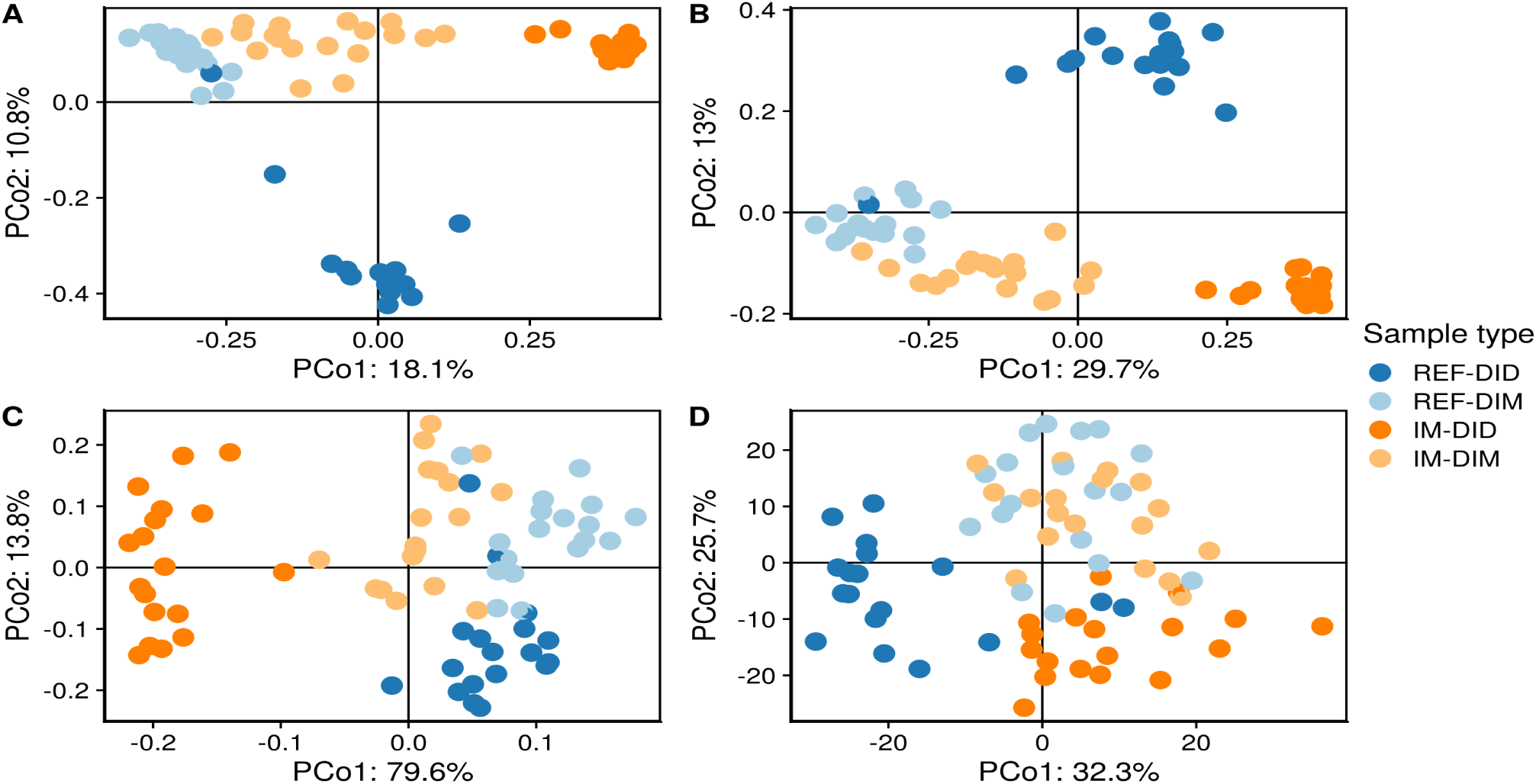
The sample origin and diet effects on the beta-diversity of distal intestinal microbiota in seawater phase Atlantic salmon. The PCoA plots were built on Jaccard (A), unweighted UniFrac (B), Aitchison (C) and phylogenetic isometric log-ratio (PHILR) transformed Euclidean (D) distance matrix, respectively. PCo, principle coordinate; REF, reference diet; IM, insect meal diet; DID, distal intestine digesta; DIM, distal intestine mucosa.

The PERMANOVA and its following conditional contrasts largely confirmed the PCoA results. Regardless of the distance matrix used, both main factors had significant effects on the beta-diversity and their interaction was significant as well (*p* < 0.05) (Table 1). Results on the tests of homogeneity of multivariate dispersions are shown in Table 2. For Jaccard distance, significant differences in the multivariate dispersions were observed between digesta and mucosa samples for both diets (REF-DID VS. REF-DIM, *p* = 0.045; IM-DID VS. IM-DIM, *p* = 0.002), and between diets for digesta samples (REF-DID VS. IM-DID, *p* = 0.002). For unweighted UniFrac distance, IM-DID showed lower multivariate dispersions than other sample types resulting in significant differences compared to REF-DID (*p* = 0.002) and IM-DIM (*p* = 0.002). For Aitchison distance, REF-DIM showed lower multivariate dispersions than other sample types resulting in significant differences compared to REF-DID (*p* = 0.046) and IM-DIM (*p* = 0.046). For PHILR transformed Euclidean distance, the differences in the multivariate dispersions among the sample types were not significant (*p* > 0.05).

**Table 1.** PERMANOVA results and subsequent conditional contrasts.

| Distance matrix | Main effects |  | Interaction | Conditional contrasts |  |  |  |
| --- | --- | --- | --- | --- | --- | --- | --- |
|  | Diet | Sample origin |  | REF-DIC VS. IM-DIC | REF-DIM VS. IM-DIM | REF-DIC VS. REF-DIM | IM-DIC VS. IM-DIM |
| Jaccard | 0.001 | 0.001 | 0.001 | 0.001 | 0.001 | 0.001 | 0.001 |
| Unweighted UniFrac | 0.001<sup>1</sup> | 0.001 | 0.001 | 0.001<sup>1</sup> | 0.001 | 0.001 | 0.001 |
| Aitchison | 0.001 | 0.003 | 0.004 | 0.002 | 0.004 | 0.004<sup>1</sup> | 0.002<sup>1</sup> |
| PHILR (Euclidean)<sup>2</sup> | 0.001 | 0.001 | 0.001 | 0.001 | 0.005 | 0.001 | 0.001 |

Abbreviations: REF, reference diet; IM, insect meal diet; DID, distal intestine digesta; DIM, distal intestine mucosa.
<sup>1</sup>Monte Carlo *p* value
<sup>2</sup>Phylogenetic isometric log-ratio transformed Euclidean distance.

**Table 2.** Test of homogeneity of multivariate dispersions among groups.

| Distance matrix | Conditional contrasts |  |  |  |
| --- | --- | --- | --- | --- |
|  | REF-DID VS. IM-DID | REF-DIM VS. IM-DIM | REF-DID VS. REF-DIM | IM-DID VS. IM-DIM |
| Jaccard | 0.002 | 0.103 | 0.015 | 0.002 |
| Unweighted UniFrac | 0.008 | 0.662 | 0.151 | 0.012 |
| Aitchison | 0.453 | 0.046 | 0.046 | 0.369 |
| PHILR (Euclidean)<sup>1</sup> | 0.240 | 0.266 | 0.240 | 0.266 |

Abbreviations: REF, reference diet; IM, insect meal diet; DID, distal intestine digesta; DIM, distal intestine mucosa.
<sup>1</sup>Phylogenetic isometric log-ratio transformed Euclidean distance.

### Significant associations between microbial clades and sample metadata

The multivariate association analysis identified 53 taxa showing significant associations with the metadata of interest (Fig. 5A). The diagnostic plots showing the raw data underlying the significant associations are shown in Fig. S3–8. Forty-seven differentially abundant taxa were identified for the sample origin effect, 45 of which, including *Bacillus*, *Enterococcus*, *Flavobacterium*, *Lactobacillus*, *Lactococcus*, *Leuconostoc*, *Mycoplasma*, *Peptostreptococcus*, *Photobacterium*, *Staphylococcus*, *Streptococcus*, *Vagococcus* and *Weissella*, showed lower relative abundances in the mucosa than the digesta (Fig. S3). In contrast, two taxa belonging to the *Spirochaetes* phylum, *B. andersonii* and unclassified *Spirochaetaceae*, were enriched in the mucosa (Fig. 5B). Thirty-six differentially abundant taxa were identified for the diet effect, 26 of which showed increased relative abundances in fish fed the IM diet (Fig. S4). Among these 26 taxa, some were enriched in both intestinal digesta and mucosa which included *Actinomyces*, unclassified *Bacillaceae*, *Bacillus*, unclassified *Beutenbergiaceae*, *Brevibacterium*, *Corynebacterium 1*, *Enterococcus*, unclassified *Lactobacillales*, *Microbacterium*, *Oceanobacillus* and unclassified *RsaHF231* (partially illustrated as Fig. 5C). For the histological scores, the relative abundance of unclassified *Sphingobacteriaceae* and unclassified *RsaHF231* were found to increase and decrease, respectively, in fish scored abnormal regarding lamina propria cellularity (lpc) in distal intestine (Fig. S5). The relative abundance of *Acinetobacter* and *Pseudomonas* were negatively correlated with the distal intestine somatic index (DISI) (Fig. S6). Six taxa, including *Actinomyces*, *B. andersonii*, *Kurthia*, *Lysobacter*, *Microbacterium* and the unclassified *Sphingobacteriaceae*, were found to associate with the expression of genes related to immune responses (Fig. S7). Notably, the relative abundance of *B. andersonii* showed a clear positive correlation with the expression levels of immune genes (Fig. 5D), which decreased as the PC1 of the PCA increased. Furthermore, 3 taxa including *Cellulosimicrobium*, *Glutamicibacter* and the unclassified *Spirochaetaceae* were found to associate with the expression of genes related to barrier functions (Fig. S8). The relative abundance of the unclassified *Spirochaetaceae* showed a negative correlation with the expression levels of barrier function relevant genes (Fig. 5E), which decreased as the PC1 of the PCA increased.

**Fig. 5.**
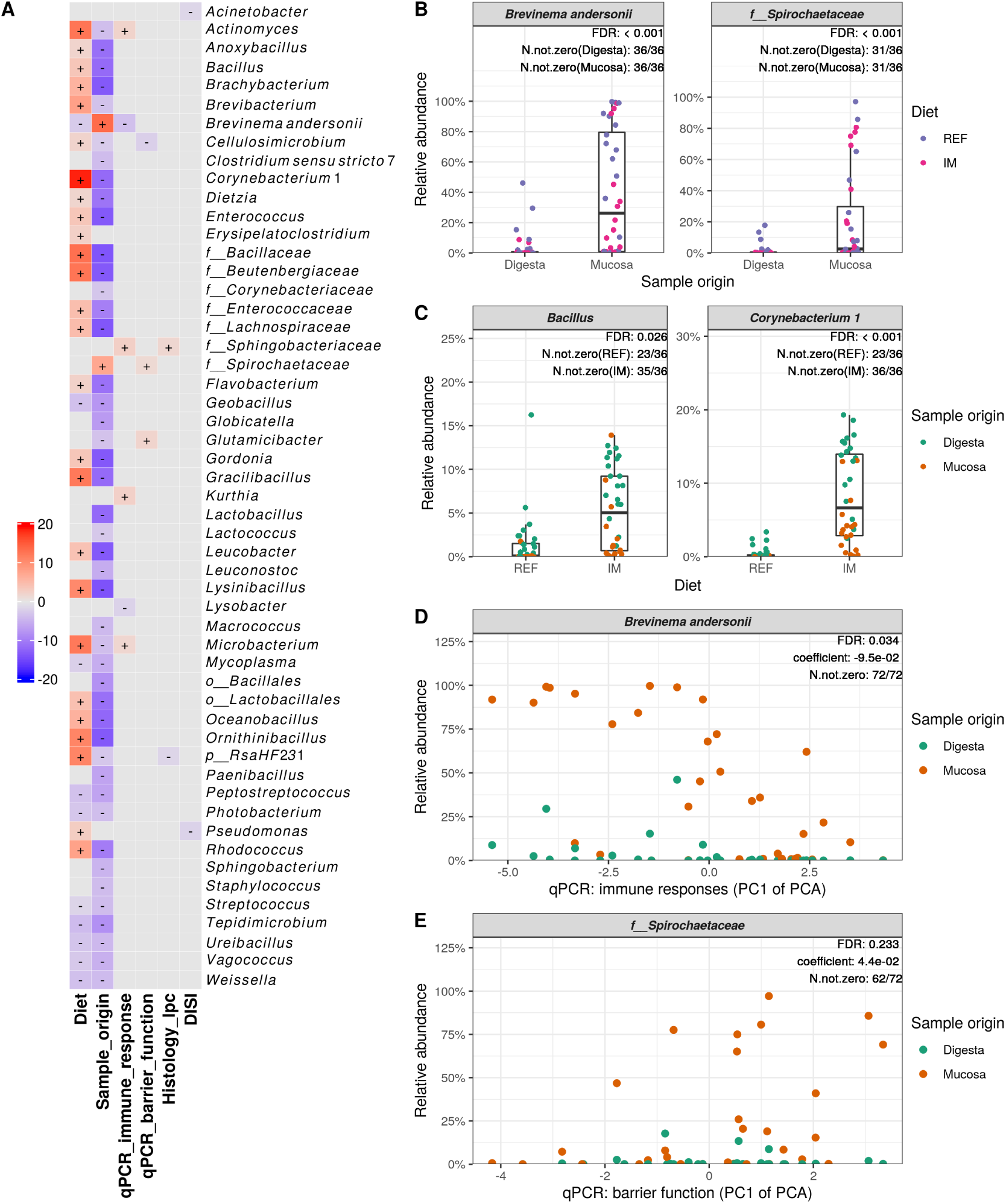
Significant associations between microbial clades and sample metadata. (A) Heatmap summarizing all the significant associations between microbial clades and sample metadata. Color key: -log(*q*-value) * sign(coefficient). Cells that denote significant associations are colored (red or blue) and overlaid with a plus (+) or minus (-) sign that indicates the direction of association: Diet (+), higher abundance in salmon fed the IM diet; Sample_origin (+), higher abundance in mucosa samples; Histology_lpc (+), higher abundance in salmon scored abnormal regarding lamina propria cellularity (lpc) in the distal intestine; DISI (+), positive correlation between microbial clade abundance and distal intestine somatic index (DISI); qPCR_immune_response (+) / qPCR_barrier_function (+), negative correlation between microbial clade abundance and the gene expression levels. (B) Taxa that are more abundant in the intestinal mucosa than the digesta. (C) Representative taxa showing increased relative abundances in both intestinal digesta and mucosa of salmon fed the IM diet. (D) The positive correlation between the relative abundance of *B. andersonii* and immune gene expression levels in the distal intestine. Note that the expression levels of the immune genes decreased as the PC1 of the PCA increased. (E) The negative correlation between the relative abundance of the unclassified *Spirochaetaceae* and the expression levels of barrier function relevant genes. Also note that the expression levels of the barrier function relevant genes decreased as the PC1 of the PCA increased. *p*_, phylum; *o*_, order; *f*_, family; FDR, false discovery rate; N.not.zero, number of observations that are not zero; REF, reference diet; IM, insect meal diet.

## Discussion

### Core microbiota

In accordance with previous studies in Atlantic salmon (20, 33–38), *Aliivibrio*, *B. andersonii* and *Mycoplasma* were identified as core microbiota in the present study. *Aliivibrio* is commonly found in the seawater phase Atlantic salmon intestine (35–37, 39–43) and has been identified as a core taxon of both wild and captive Atlantic salmon (34, 36, 37). Provided its common presence in seawater, *Aliivibrio* may have originated from the surrounding water and colonized the intestinal mucosa as Atlantic salmon constantly drink seawater to prevent dehydration in a hyperosmotic environment. Currently, the taxon *Aliivibrio* comprises of four closely related species including *Aliivibrio fischeri*, *Aliivibrio logei*, *Aliivibrio salmonicida* and *Aliivibrio wodanis*, which were split from the *Vibrio* genus and reclassified as *Aliivibrio* in 2007 (44). Strains of *A. fischeri* and *A. logei* have been described as bioluminescent symbionts of certain fishes and squids (45), whereas *A. salmonicida* and *A. wodanis* have been identified as pathogens for Atlantic salmon causing cold-water vibriosis (46) and ‘winter ulcer’ (47), respectively.

Though *Spirochaetes* has typically been found in low abundances in the Atlantic salmon intestine (19, 23, 27, 39), two recent studies have identified *B. andersonii* as a core taxon of both digesta- and mucosa-associated intestinal microbiota in seawater phase Atlantic salmon (35, 36). Furthermore, *B. andersonii* is also a predominant taxon in the digesta and mucosa in one of the studies (36). *B. andersonii* was initially isolated from short-tailed shrews (*Blarina brevicauda*) and white-footed mice (*Peromyscus leucopus*) as an infectious pathogen (48). This taxon has also been found in the intestine and gill tissue of rainbow trout (*Oncorhynchus mykiss*) (49), and intestinal digesta of Senegalese sole (*Solea senegalensis*)(50).

*Mycoplasma* is widely distributed in nature and well known for its minute size and lack of cell wall. Like *Aliivibrio*, *Mycoplasma* has been frequently identified as a core taxon of both wild and captive Atlantic salmon as well (20, 33, 35–38). It was found to be more abundant in marine adults than in freshwater juvenile Atlantic salmon (37) and sporadically predominate intestinal microbial community in the digesta (20, 36, 37, 41, 51) and mucosa (35) reaching as high as > 90% of total read counts in extreme cases. Due to its small compact genome and limited biosynthesis capacities, *Mycoplasma* typically forms obligate parasitic or commensal relationships with its host to obtain necessary nutrients such as amino acids, fatty acids and sterols (52). The major acids produced by *Mycoplasma* during fermentation are lactic acid and acetic acid (53), the latter of which was found in our recent studies to be 2-4 orders of magnitude higher than other short-chain fatty acids in the Atlantic salmon distal intestine (unpublished data). A recent study found that Atlantic salmon *Mycoplasma*, recovered by shotgun metagenomic sequencing, was closely related to *Mycoplasma penetrans* and *Mycoplasma yeatsii* (20). Compared to these closely related *Mycoplasma* species, the salmon *Mycoplasma* appears to carry a much lower number of unique genes that are enriched in carbohydrate uptake and low in peptidase synthesis.

### Sample origin effect

In line with previous findings in mammals and fish (19, 22–25), we observed substantial differences between digesta- and mucosa-associated microbiota. The microbial richness and diversity were much higher in the digesta than the mucosa, as previously observed in seawater phase Atlantic salmon (19, 23, 26). Furthermore, most of the bacterial taxa in the distal intestine, including those commonly found in the Atlantic salmon intestine such as *Bacillus*, *Enterococcus*, *Flavobacterium*, *Lactobacillus*, *Lactococcus*, *Leuconostoc*, *Mycoplasma*, *Peptostreptococcus*, *Photobacterium*, *Staphylococcus*, *Streptococcus*, *Vagococcus* and *Weissella*, were less abundant in the mucosa than in the digesta. These results are suggestive of a selection pressure from the host that determines which microbial clades colonize and flourish in the intestinal mucus layer (54). In this study, two taxa belonging to the *Spirochaetes* phylum, *B. andersonii* and unclassified *Spirochaetaceae*, were more abundant in the distal intestine mucosa than the digesta. As aforementioned, *Spirochaetes* were typically found in low abundances in the Atlantic salmon intestine. Yet a recent study also showed that irrespective of diets *B. andersonii* seemed to be more abundant in the intestinal mucosa than the digesta of seawater phase Atlantic salmon (36). Known for high motility and chemotactic attraction to mucin, some *Spirochaetes* can penetrate the mucus and associate with the intestinal mucosa (55–57). Further work is required to confirm whether these taxa are consistently enriched in the intestinal mucus layer of seawater phase Atlantic salmon.

### Diet effect

Diet is one of the key factors in shaping the fish intestinal microbiota. In agreement with previous findings in rainbow trout (29, 58, 59) and laying hens (60, 61), we found that the insect meal diet altered the distal intestinal microbiota assemblage resulting in higher microbial richness and diversity. Our findings, showing that the insect meal diet increased the relative abundance of *Actinomyces*, *Bacillus*, *Brevibacterium*, *Corynebacterium 1* and *Enterococcus*, are in accord with recent studies in rainbow trout fed diets containing 30% black soldier fly larvae meal (29, 59). Importantly, these results were partly confirmed in other studies employing fluorescence *in situ* hybridization for targeted profiling of changes in the intestinal microbiota. Specifically, increased absolute abundance of *Lactobacillus*/*Enterococcus* was found in rainbow trout fed 20% dietary black soldier fly larvae meal (62), whereas increased absolute abundance of *Bacillus*, *Enterococcus* and *Lactobacillus* was documented in Siberian sturgeon (*Acipenser baerii*) fed 15% black soldier fly larvae meal (63).

The increases in the relative abundance of specific microbial clades in Atlantic salmon fed the insect meal diet may be explained by feed-borne microbiota and/or feed composition. Bacterial taxa, including *Actinomyces*, *Bacillus*, *Brevibacterium*, *Corynebacterium*, *Enterococcus*, *Oceanobacillus* and *RsaHF231*, have been found in black soldier fly whole larvae or larvae intestine (64–67). The fact that *RsaHF231* has not been documented in fish before indicates that these bacterial taxa may have partially originated from black soldier fly larvae meal. Our results from the freshwater feeding trial showed that these bacterial taxa were also enriched in the intestinal digesta and mucosa of Atlantic salmon smolts fed an insect meal diet containing 60% soldier fly larvae meal. Importantly, these bacterial taxa were also detected in the feed pellets which contained considerable amount of bacterial DNA that is comparable to intestinal digesta (unpublished data). Given the hydrothermal treatments the feed pellets underwent during the extrusion, the feed-borne microbiota profiled by the DNA sequencing techniques could have largely originated from dead bacteria and bacterial spores rather than living bacteria. As sequencing-based methods cannot differentiate between living and dead cells, future studies should investigate to what extent the feed-borne microbiota may contribute to, or confound the observed diet effects on intestinal microbiota, using methods that distinguish living and dead bacteria such as viability PCR and RNA sequencing (68). On the other hand, unique nutrients in the insect meal diet such as chitin, an essential component of the insect exoskeleton, may have selectively promoted the growth of certain intestinal microbes. *Actinomyces* species are often identified as active chitin degraders showing enhanced growth and activity upon chitin addition (69). Many bacterial species belonging to *Bacillus* can produce chitinase (70). *Bacillus* and *Lactobacillus* were two of the predominant taxa in the intestinal mucosa of Atlantic salmon fed a 5% chitin diet, the former of which displayed the highest *in vitro* chitinase activity (71).

### Significant interactions between diet and sample origin effect

We observed in the present study that the diet effect on the intestinal microbial community richness and structure was dependent on the sample origin, with mucosa-associated intestinal microbiota showing higher resilience to the dietary change. Our results corroborate previous findings in rainbow trout revealing that mucosa-associated intestinal microbiota was less influenced by dietary inclusion of 30% black soldier fly larvae meal compared to digesta-associated intestinal microbiota (28, 29). Results from molecular-based studies on salmonid intestinal microbiota hitherto suggest that diet modulates digesta- and mucosa-associated intestinal microbiota to varying degrees with the latter generally being more resilient to dietary interventions (19, 26–29, 35). As such, current practices of profiling only one of or a mixture of digesta- and mucosa-associated microbiota may obscure the response of intestinal microbiota to dietary changes. To fully unveil the response of intestinal microbiota to dietary changes, we recommend concurrent profiling of digesta- and mucosa-associated intestinal microbiota whenever it is feasible.

### Significant associations between microbial clades and sample metadata

To our knowledge, only a few studies have carried out association analysis between intestinal microbial clades and host responses in Atlantic salmon. As such, our results should be treated as preliminary observations and critically evaluated in later studies. Herein, we highlight the significant associations between two mucosa-enriched taxa and host gene expressions in the intestine. Specifically, *B. andersonii*, part of the core microbiota, was associated with the expression of genes related to pro- and anti-inflammatory responses whereas the unclassified *Spirochaetaceae* was associated with the expression of genes related to barrier function. Intestinal microbiota is well known to modulate the local immune responses and intestinal epithelial barrier function (72). Furthermore, it is hypothesized that mucosa-associated microbiota plays a more crucial role in shaping the host immunity in that it can interact both directly and indirectly with intestinal epithelial barrier whereas digesta-associated microbiota can only interact indirectly (54). Taken together, further research should be undertaken to investigate the potential ecological and functional significance of these two taxa for seawater phase Atlantic salmon.

### Quality control: use of mock and negative controls

As in any field of research, conducting a well-controlled microbiome study requires great care in the experiment design such as setting up appropriate experimental controls. The use of mock as a positive control allows for critical evaluation and optimization of microbiota profiling workflow. That all the bacterial taxa in the mock were correctly identified at the genus level indicates that the current workflow is reliable for the taxonomic profiling of intestinal microbiota. Furthermore, the taxonomic profile of mock from different DNA extraction batches was fairly similar, suggesting that the results generated by the current workflow are also largely reproducible. However, the low concordance between the expected and observed relative abundance of bacterial taxa in the mock is reminiscent of the fact that bias is introduced at different steps of the marker-gene survey (73–75), among which DNA extraction and PCR amplification are the two largest sources of bias due to preferential extraction and amplification of some microbial clades over others. In line with previous observations that Gram-positive bacteria may be more subjective to incomplete lysis during DNA extraction due to their tough cell walls (76, 77), the recovery of most Grampositive bacteria in the mock was lower than the expected. The insufficient lysing of Gram-positive bacteria in the mock was largely mitigated in our later experiments by using a mixture of beads with different sizes for the bead beating during DNA extraction (unpublished data). The bias in the markergene sequencing experiments, as reflected in the observed taxonomic profile of the mock, highlights the necessity of validating such results by absolute quantification techniques such as cultivation (if possible), qPCR, flow cytometry and fluorescence *in situ* hybridization.

Reagent contamination is a common issue in molecular-based studies of microbial communities. The main contaminating taxon identified in this study is *Pseudomonas*, which has been reported as a common reagent contaminant in numerous studies (78–84). Given the dominance of *Pseudomonas* in the negative controls of both DNA extraction and PCR, most of the observed contamination has likely derived from PCR reagents such as molecular-grade water (85–87). Notably, *Pseudomonas* has also been isolated from intestinal digesta and mucosa of Atlantic salmon by traditional culturing approaches (71, 88–90), and reported as a member of Atlantic salmon core microbiota in culture-independent studies (19, 23, 33, 34, 38, 91). Due to the low taxonomic resolution of amplicon sequencing, it is difficult to discern contaminating taxa from true signals solely based on taxonomic labels. The inclusion of negative controls, coupled with quantifications of microbial DNA concentration in the samples, has enabled fast and reliable identification of contaminating taxa in this study. Besides *Pseudomonas*, other common reagent contaminants, including *Bradyrhizobium*, *Burkholderia*, *Comamonas*, *Methylobacterium*, *Propionibacterium*, *Ralstonia*, *Sphingomonas* and *Stenotrophomonas* (83, 85, 87, 92–96), have also been frequently reported as members of Atlantic salmon intestinal microbiota, indicating that existing studies of Atlantic salmon intestinal microbiota may have been plagued with reagent contamination that is hard to ascertain due to lack of negative controls. As reagent contamination is unavoidable, study-specific and can critically influence sequencing-based microbiome analyses (85, 97, 98), negative controls should always be included and sequenced in microbiome studies especially when dealing with low microbial biomass samples like intestinal mucosa.

### Conclusion

In summary, we confirmed previous findings in mammals and fish that intestinal digesta and mucosa harbor microbial communities with clear differences. Regardless of diet, microbial richness and diversity were much higher in the digesta than the mucosa. The insect meal diet altered the distal intestinal microbiota assemblage resulting in higher microbial richness and diversity. The diet effect was however dependent on the sample origin, with mucosa-associated intestinal microbiota being more resilient to the dietary change. To fully unveil the response of intestinal microbiota to dietary changes, concurrent profiling of digesta- and mucosa-associated intestinal microbiota is recommended whenever feasible. Lastly, we identified two mucosa-enriched taxa, *Brevinema andersonii* and unclassified *Spirochaetaceae*, which seemed to be associated with the expression in the distal intestine of genes related to immune responses and barrier function, respectively. As mucosa-associated microbiota could play a more critical role in shaping the host metabolism, their potential functional significance for seawater phase Atlantic salmon merits further investigations.

## Materials and Methods

### Experimental fish, diet and sampling

A 16-week seawater feeding trial with Atlantic salmon (initial body weight = 1.40 kg, S.D. = 0.043 kg) was conducted at the Gildeskål Research Station (GIFAS), Nordland, Norway, in accordance with laws regulating the experimentation with live animals in Norway. The experimental fish were randomly assigned into 6 net pens (5 × 5 × 5 m) each containing 90 fish. The fish were fed, in triplicate net pens, either a commercially-relevant reference diet (REF) with a combination of fish meal, soy protein concentrate, pea protein concentrate, corn gluten and wheat gluten as the protein source, or an insect meal diet (IM) wherein all the fish meal and most of the pea protein concentrate were replaced by black soldier fly larvae meal. Fish were fed by hand until apparent satiation once or twice daily depending on the duration of daylight. During the feeding trial, the water temperature ranged from 7 °C to 13 °C. Further details on the formulation and chemical composition of the diets, and insect meal have been reported elsewhere (31, 32).

At the termination of the feeding trial, 6 fish were randomly taken from each net pen, anesthetized with tricaine methanesulfonate (MS222®; Argent Chemical Laboratories, Redmond, WA, USA) and euthanized by a sharp blow to the head. After cleaning the exterior of each fish with 70% ethanol, the distal intestine, i.e., the segment from the increase in intestinal diameter and the appearance of transverse luminal folds to the anus, was aseptically removed from the abdominal cavity, placed in a sterile Petri dish and opened longitudinally. Only fish with digesta along the whole intestine were sampled to ensure that the intestine had been exposed to the diets. The intestinal digesta was collected into a 50 mL skirted sterile centrifuge tube and mixed thoroughly using a spatula. An aliquot of the homogenate was then transferred into a 1.5 mL sterile Eppendorf tube and snap-frozen in liquid N_2_ for the profiling of digesta-associated intestinal microbiota. A tissue section from the mid part of the distal intestine was excised and rinsed in sterile phosphate-buffered saline 3 times to remove traces of the remaining digesta. After rinsing, the intestinal tissue was longitudinally cut into 3 pieces for histological evaluation (fixed in 4% phosphate-buffered formaldehyde solution for 24 h and transferred to 70% ethanol for storage), RNA extraction (preserved in RNAlater solution and stored at −20 °C) and profiling of mucosa-associated intestinal microbiota (snap-frozen in liquid N_2_), respectively. The collection of microbiota samples was performed near a gas burner to secure aseptic conditions. After the sampling of each fish, tools were cleaned and decontaminated by a 70% ethanol spray and flaming. Microbiota samples of the distal intestine digesta (DID) and mucosa (DIM) were transported in dry ice and stored at −80 °C until DNA extraction.

### DNA extraction

Total DNA was extracted from ~200 mg distal intestine digesta or mucosa using the QIAamp® DNA Stool Mini Kit (Qiagen, Hilden, Germany; catalog no., 51504) with some modifications to the manufacturer’s specifications as described before (19), except that 2 mL prefilled bead tubes (Qiagen; catalog no., 13118-50) were used for the bead beating. For quality control purposes, a companion “blank extraction” sample was added to each batch of sample DNA extraction by omitting the input material, whereas an additional microbial community standard (Zymo-BIOMICS™, Zymo Research, California, USA; catalog no., D6300), i.e. mock, was included for each DNA extraction kit as a positive control. The mock consists of 8 bacteria (*Pseudomonas aeruginosa*, *Escherichia coli*, *Salmonella enterica*, *Lactobacillus fermentum*, *Enterococcus faecalis*, *Staphylococcus aureus*, *Listeria monocytogenes*, *Bacillus subtilis*) and 2 yeasts (*Saccharomyces cerevisiae*, *Cryptococcus neoformans*).

### Amplicon PCR

The V1-2 hypervariable regions of the bacterial 16S rRNA gene were amplified using the primer set 27F (5’-AGA GTT TGA TCM TGG CTC AG-3’) and 338R (5’- GCW GCC WCC CGT AGG WGT-3’) (99). The PCR was run in a total reaction volume of 25 µL containing 12.5 µL of Phusion® High-Fidelity PCR Master Mix (Thermo Scientific, CA, USA; catalog no., F531L), 10.9 µL molecular grade H_2_O, 1 µL DNA template and 0.3 µL of each primer (10 µM). The amplification program was set as follows: initial denaturation at 98 °C for 3 min; 35 cycles of denaturation at 98 °C for 15 s, annealing decreasing from 63 °C to 53 °C in 10 cycles for 30 s followed by 25 cycles at 53 °C for 30 s, and extension at 72 °C for 30 s; followed by a final extension at 72 °C for 10 min. For samples with faint or invisible bands in the agarose gel after PCR, the PCR condition was optimized by applying serial dilutions to the DNA templates to reduce the influence of PCR inhibitors. All the digesta samples were diluted 1:2 in buffer ATE (10 mM Tris-Cl, pH 8.3, with 0.1 mM EDTA and 0.04% NaN_3_) whereas all the mucosa samples were diluted 1:32. The formal amplicon PCR was run in duplicate incorporating two negative PCR controls, which were generated by replacing the template DNA with molecular grade H_2_O. The duplicate PCR products were then pooled and examined by a 1.5% agarose gel electrophoresis.

### Quantification of 16S rRNA gene by qPCR

To assist in identifying contaminating sequences, the 16S rRNA gene quantity in the diluted DNA templates used for the amplicon PCR was measured by qPCR. The qPCR assays were performed using a universal primer set (forward, 5’-CCA TGA AGT CGG AAT CGC TAG-3’; reverse, 5’-GCT TGA CGGGCG GTG T-3’) that has been used for bacterial DNA quantification in previous studies (100, 101). The assays were carried out using the LightCycler 96 (Roche Applied Science, Basel, Switzerland) in a 10 µL reaction volume, which contained 2 µL of PCR-grade H_2_O, 1 µL diluted DNA template, 5 µL LightCycler 480 SYBR Green I Master Mix (Roche Applied Science) and 1 µL (3 µM) of each primer. Samples, together with the extraction blanks and mock, were run in duplicate in addition to Femto™ bacterial DNA standards (Zymo Research; catalog no., E2006) and a no-template control of the qPCR assay. The qPCR program encompassed an initial enzyme activation step at 95 °C for 2 min, 45 three-step cycles of 95 °C for 10 s, 60 °C for 30 s and 72 °C for 15 s, and a melting curve analysis at the end. Quantification cycle (Cq) values were determined using the second derivative method (102). The specificity of qPCR amplification was confirmed by evaluating the melting curve of qPCR products and the band pattern on the agarose gel after electrophoresis. The inter-plate calibration factor was calculated following the method described in (103), using the bacterial DNA standards as inter-plate calibrators.

### Sequencing

The sequencing was carried out on a Miseq platform following the Illumina 16S metagenomic sequencing library preparation protocol (104). Briefly, the PCR products were cleaned using the Agencourt AMPure XP system (Beckman Coulter, Indiana, USA; catalog no., A63881), multiplexed by dual indexing using the Nextera XT Index Kit (Illumina, California, USA; catalog no., FC-131-1096) and purified again using the AMPure beads. After the second clean-up, representative libraries were selected and analyzed using the Agilent DNA 1000 Kit (Agilent Technologies, California, USA; catalog no., 5067-1505) to verify the library size. Cleaned libraries were quantified using the Invitrogen Qubit™ dsDNA HS Assay Kit (Thermo Fisher Scientific, California, USA; catalog no., Q32854), diluted to 4 nM in 10 mM Tris (pH 8.5) and finally pooled in an equal volume. Negative controls with library concentrations lower than 4 nM were pooled in equal volume directly. Due to the low diversity of amplicon library, 15% Illumina generated PhiX control (catalog no., FC-110-3001) was spiked in by combining 510 µL amplicon library with 90 µL PhiX control library. The library was loaded at 6 pM and sequenced using the Miseq Reagent Kit v3 (600-cycle) (Illumina; catalog no., MS-102-3003).

### Sequence data processing

The raw sequence data were processed by the DADA2 1.14 in R 3.6.3 (105) to infer amplicon sequence variants (ASVs) (106). Specifically, the demultiplexed paired-ended reads were trimmed off the primer sequences (forward reads, first 20 bps; reverse reads, first 18 bps), truncated at the position where the median Phred quality score crashed (forward reads, at position 290 bp; reverse reads, at position 248 bp) and filtered off low-quality reads. After trimming and filtering, the run-specific error rates were estimated and the ASVs were inferred by pooling reads from all the samples sequenced in the same run. The chimeras were removed using the “pooled” method after merging the reads. The resulting raw ASV table and representative sequences were imported into QIIME2 (version, 2020.2) (107). The taxonomy was assigned by a scikitlearn naive Bayes machine-learning classifier (108), which was trained on the SILVA 132 99% OTUs (109) that were trimmed to only include the regions of 16S rRNA gene amplified by our primers. Taxa identified as chloroplasts or mitochondria were excluded from the ASV table. The ASV table was conservatively filtered to remove ASVs that had no phylum-level taxonomic assignment or appeared in only one biological sample. Contaminating ASVs were identified based on two suggested criteria: contaminants are often found in negative controls and inversely correlate with sample DNA concentration (84). The ASVs filtered from the raw ASV table were also removed from the representative sequences, which were then inserted into a reference phylogenetic tree built on the SILVA 128 database using SEPP (110). The alpha rarefaction curves and the core metrics results were generated with a sampling depth of 10000 and 2047 sequences per sample, respectively. For downstream data analysis and visualization, QIIME2 artifacts were imported into R using the qiime2R package (111) and a phyloseq (112) object was assembled from the sample metadata, ASV table, taxonomy and phylogenetic tree. The core microbiota and alpha-diversity indices were computed using the ASV table collapsed at the species level. The core microbiota was calculated based on the 80% prevalence threshold and visualized by the Venn’s diagram. The alpha-diversity indices, including observed species, Pielou’s evenness, Shannon’s index and Faith’s phylogenetic diversity (PD), were computed via the R packages microbiome (113) and picante (114). For beta-diversity analyses, we used distance matrices including Jaccard distance, unweighted UniFrac distance, Aitchison distance and phylogenetic isometric log-ratio (PHILR) transformed Euclidean distance. Since rarefying remains to be the best solution for unweighted distance matrices (115), the Jaccard distance and unweighted UniFrac distance were computed in QIIME2 using the rarefied ASV table. The compositionality-aware distance matrices, Aitchison distance and PHILR transformed Euclidean distance, were calculated using the unrarefied ASV table. The Aitchison distance was computed by the DEICODE plugin in QIIME2, a form of Aitchison distance that is robust to high levels of sparsity by using the matrix completion to handle the excessive zeros in the microbiome data (116). The PHILR transform of the ASV table was performed in R using the philr package (117). The selected distance matrices were explored and visualized by the principal coordinates analysis (PCoA).

### Multivariate association analysis

The ASV table was collapsed at the genus level before running the multivariate association analysis. Bacterial taxa of very low abundance (< 0.01%) or low prevalence (present in < 25% of samples) were removed from the feature table. The microbial clades were then tested for significant associations with metadata of interest by MaAsLin2 (version, 0.99.12) (https://huttenhower.sph.harvard.edu/maaslin2) in R, using the default parameters. The results of the analysis are the associations of specific microbial clades with metadata, deconfounding the influence of other factors included in the model. Association was considered significant when the *q*-value was below 0.25. Metadata included in the multivariate association testing are fixed factors Diet + Sample origin + distal intestine somatic index (DISI) + lamina propria cellularity (histological scores) + immune response (qPCR) + barrier function (qPCR), and random factors FishID + NetPen. FishID was nested in NetPen, and NetPen nested in Diet. Lamina propria cellularity reflects the severity of inflammation in the distal intestine. Based on the degree of cellular infiltration within the lamina propria, a value of normal, mild, moderate, marked or severe was assigned. To make the data appropriate for the association testing, the highly skewed five-category scores were collapsed into more balanced binary data, i.e., normal and abnormal. The immune-related genes included for the association testing were myeloid differentiation factor 88 (*myd88*), interleukin 1β (*il1β*), interleukin 8 (*il8*), cluster of differentiation 3 γδ (*cd3γδ*), transforming growth factor β1 (*tgfβ1*), interferon γ (*ifnγ*), interleukin 17A (*il17a*), fork-head box P3 (*foxp3*) and interleukin 10 (*il10*), whose expression levels were higher in the distal intestine of fish assigned abnormal regarding lamina propria cellularity. Since the expression levels of immune-related genes were highly correlated, we ran a principal component analysis (PCA) and extracted the first principle component (PC1) for the association testing to avoid multicollinearity and reduce the number of association testing. For genes relevant to the barrier function, which included claudin-15 (*cldn15*), claudin-25b (*cldn25b*), zonula occludens 1 (*zo1*), E-cadherin / cadherin 1 (*cdh1*) and mucin-2 (*muc2*), we also used the PC1 of the PCA for the association testing based on the same considerations.

### Statistics

All the statistical analyses were run in R except for the PERMANOVA, which was run in PRIMER v7. The differences in the alpha-diversity indices were compared using linear mixed-effects models via the lme4 package (118). Predictor variables in the models included the fixed effects Diet + Sample origin + Diet × Sample origin, and the random effects FishID + NetPen. The models were validated by visual inspections of residual diagnostic plots generated by the ggResidpanel package (119). The statistical significance of fixed predictors was estimated by Type III ANOVA with Kenward-Roger’s approximation of denominator degrees of freedom via the lmerTest package (120). When the interaction between the main effects was significant, conditional contrasts for the main effects were made via the emmeans package (121). To compare the differences in beta-diversity, we performed the PERMANOVA (122) using the same predictors included in the linear mixed-effects models. Terms with negative estimates for components of variation were sequentially removed from the model via term pooling, starting with the one showing the smallest mean squares. At each step, the model was reassessed whether more terms needed to be removed or not. Conditional contrasts for the main effects were constructed when their interaction was significant. Monte Carlo *p* values were computed as well when the unique permutations for the terms in the PERMANOVA were small (< 100). The homogeneity of multivariate dispersions among groups was visually assessed with boxplots and was formally tested by the permutation test, PERMDISP (123), via the R package vegan (124). Multiple comparisons were adjusted by the Benjamini-Hochberg procedure where applicable. Differences were regarded as significant when *p* < 0.05.

### Code availability

All the code for reproducing the results are available from the GitHub repository (https://github.com/yanxianl/Li_AqFl2-Microbiota_ASM_2020).

### Data availability

Raw sequence data are deposited at the NCBI SRA database (https://www.ncbi.nlm.nih.gov/sra) under the BioProject PRJNA555355. Other raw data and sample metadata are available from the GitHub repository (https://github.com/yanxianl/Li_AqFl2-Microbiota_ASM_2020).

## Acknowledgments

Y.L. was granted a scholarship from the China Scholarship Council to pursue his PhD degree at Norwegian University of Life Sciences. This work was a spin-off of the “AquaFly” project (grant number, 238997), funded by the Research Council of Norway and managed by the Institute of Marine Research. Costs related to this study were covered by Norwegian University of Life Sciences. The funding agencies had no role in study design, data collection and interpretation, decision to publish or preparation of the manuscript.

The authors wish to thank Ellen K. Hage for organizing the sampling and technicians at the GIFAS for their committed animal care and supports during the sampling.

T.M.K. and Å.K. concepted and designed the study. Y.L., L.B. and K.G. participated in the sample collection. Y.L., L.B. and A.J.-T. carried out the laboratory works. Y.L. performed the bioinformatics, statistical analyses and data visualization. Y.L. and L.B. completed the first draft of the manuscript. All the authors read, revised and approved the final version of the manuscript.

We declare no conflicts of interest.

## Supplementary Note 1: Supplemental tables

The supplemental tables are available on bioRXiv:

- Table S1. Contaminating features removed from the ASV table.
- Table S2. The prevalence of core taxa in different sample types.

---

**Fig. S1.**
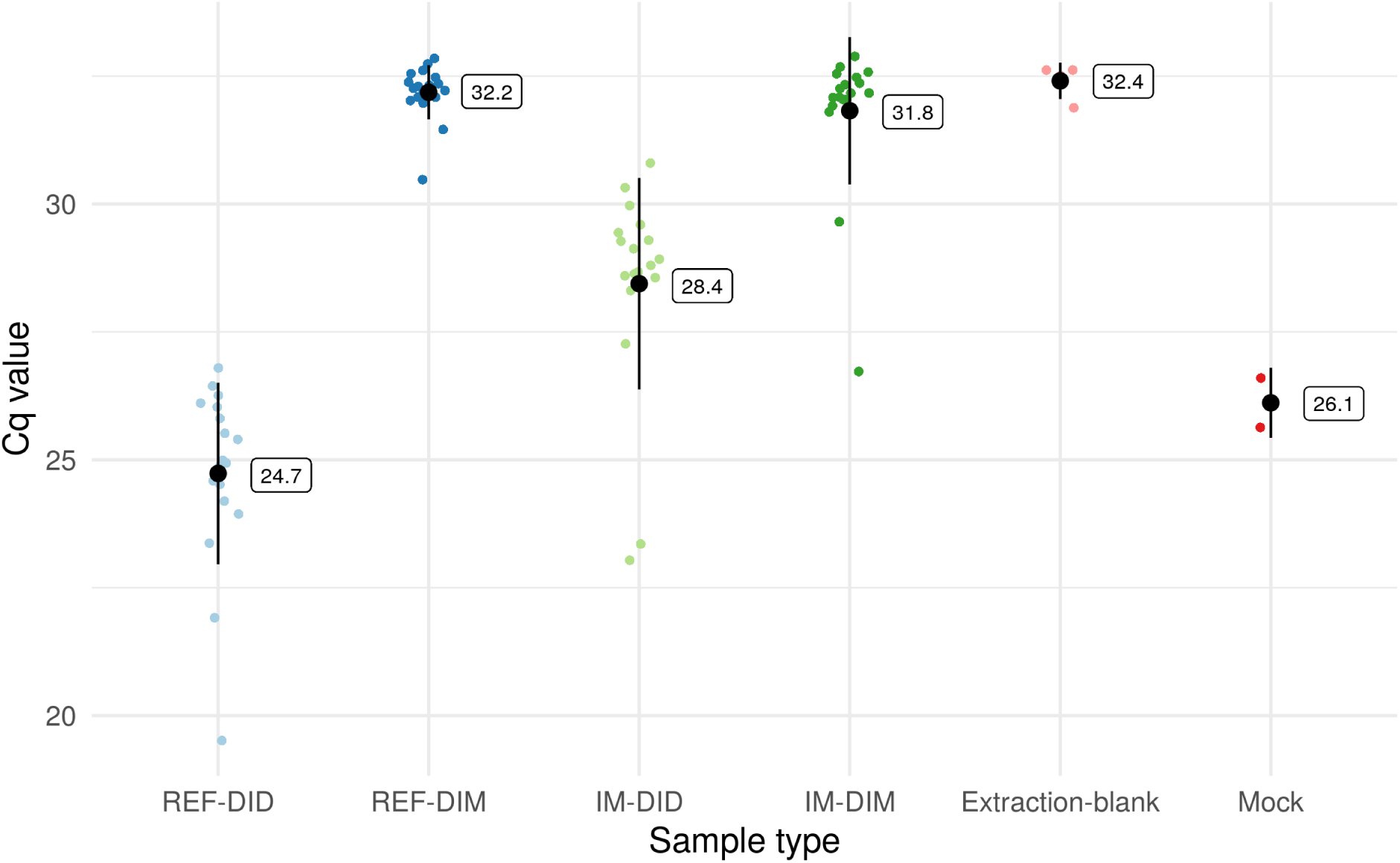
Quantification of bacterial 16S rRNA gene in different sample types using qPCR. Since the Cq values of most mucosa-associated samples were out of the linear range of the standard curve, the Cq value was used as a proxy of 16S rRNA gene quantity which is reliable for the screening of contaminant sequences. Data are presented as mean ± 1 standard deviation overlaying the raw data points. Abbreviations: REF, reference diet; IM, insect meal diet; DID, distal intestine digesta; DIM, distal intestine mucosa.

**Fig. S2.**
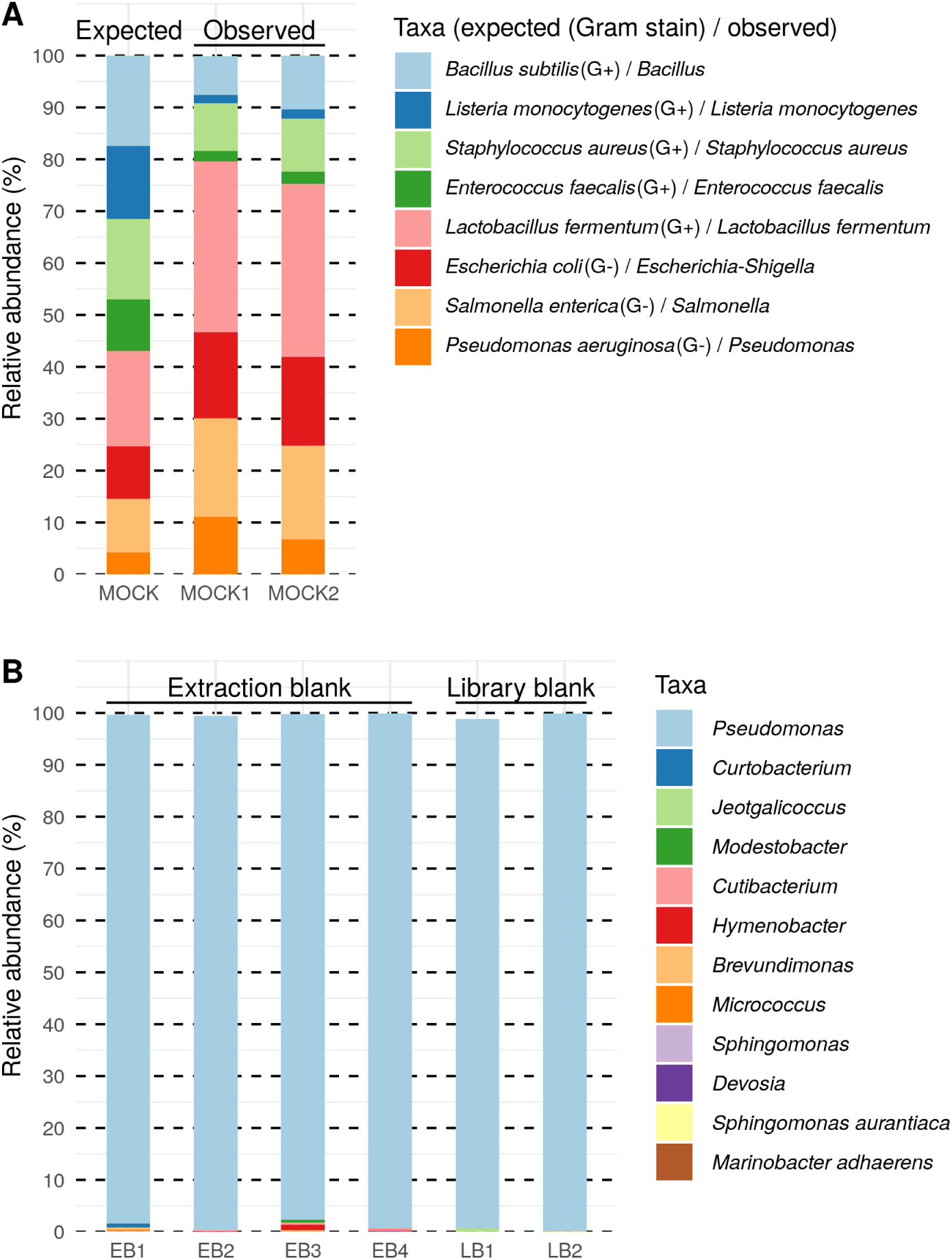
Taxonomic profile of the mock (A) and contaminating features in the negative controls (B). The lowest level of taxonomic ranks was displayed for each taxon. EB, extraction blank; LB, library blank.

**Fig. S3.**
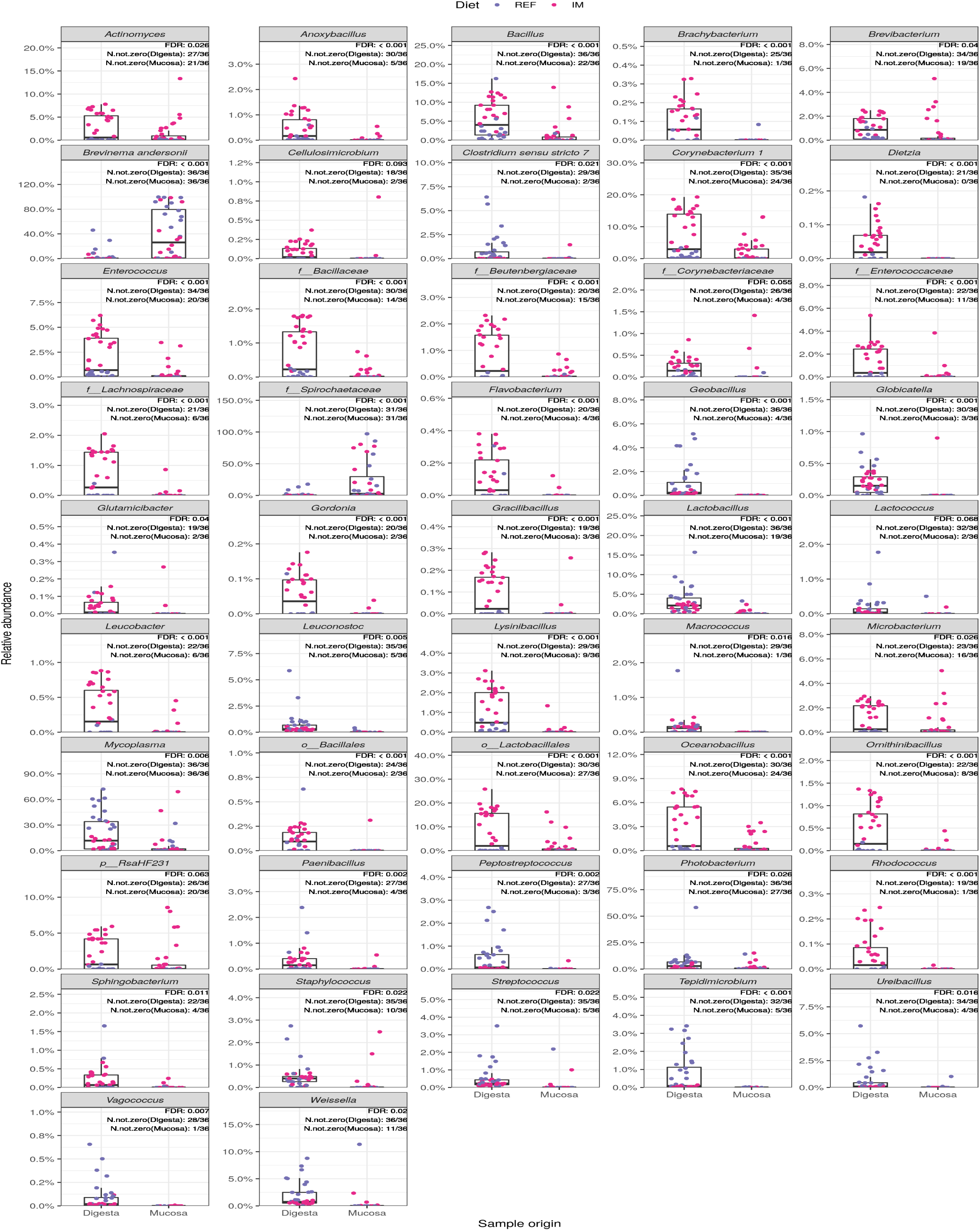
Microbial clades showing significant associations with sample origin. *p*_, phylum; *o*_, order; *f*_, family; FDR, false discovery rate; N.not.zero, number of non-zero obsetarvations; REF, reference diet; IM, insect meal diet.

**Fig. S4.**
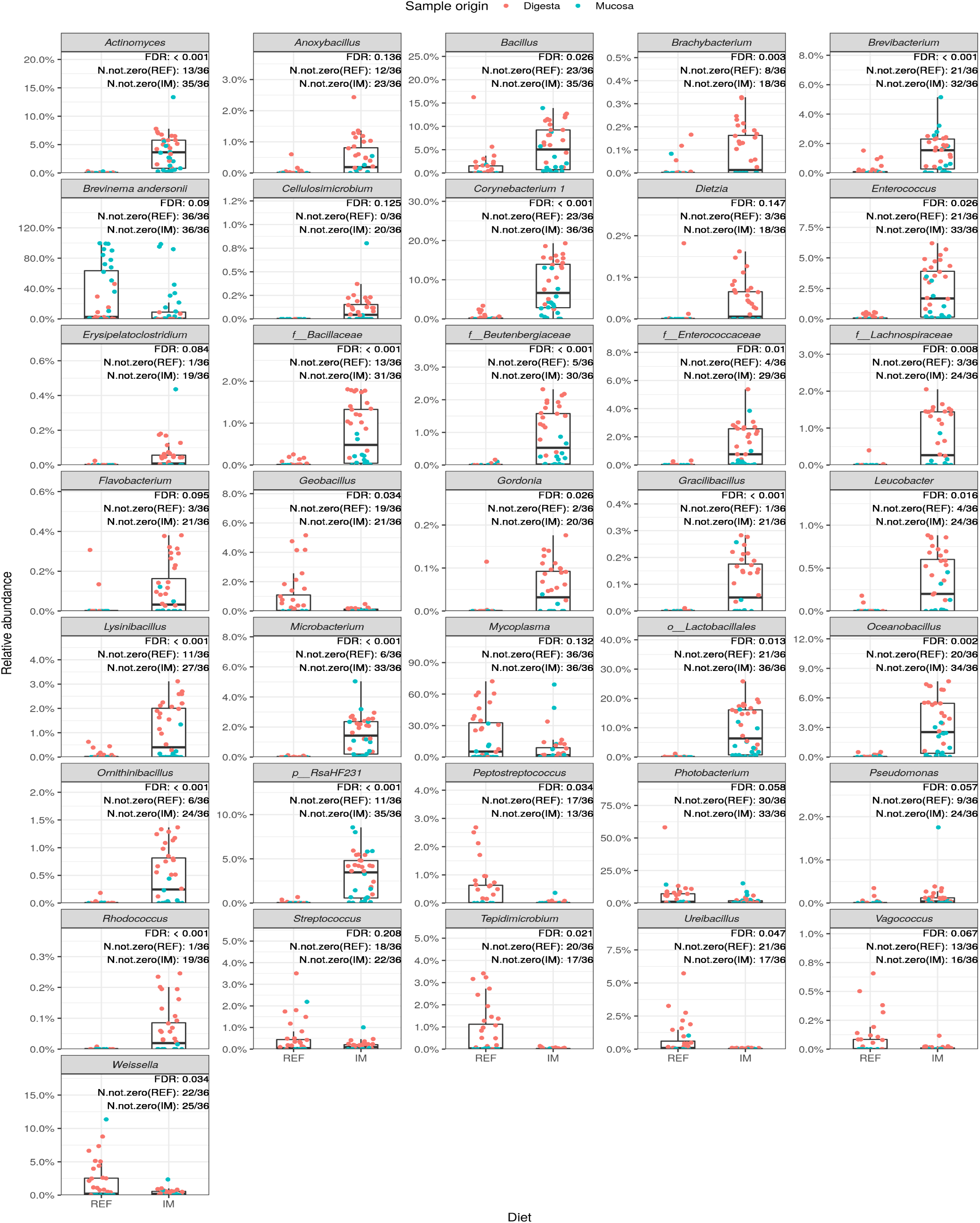
Microbial clades showing significant associations with diet. *p*_, phylum; *o*_, order; *f*_, family; FDR, false discovery rate; N.not.zero, number of non-zero observations; REF, reference diet; IM, insect meal diet.

**Fig. S5.**
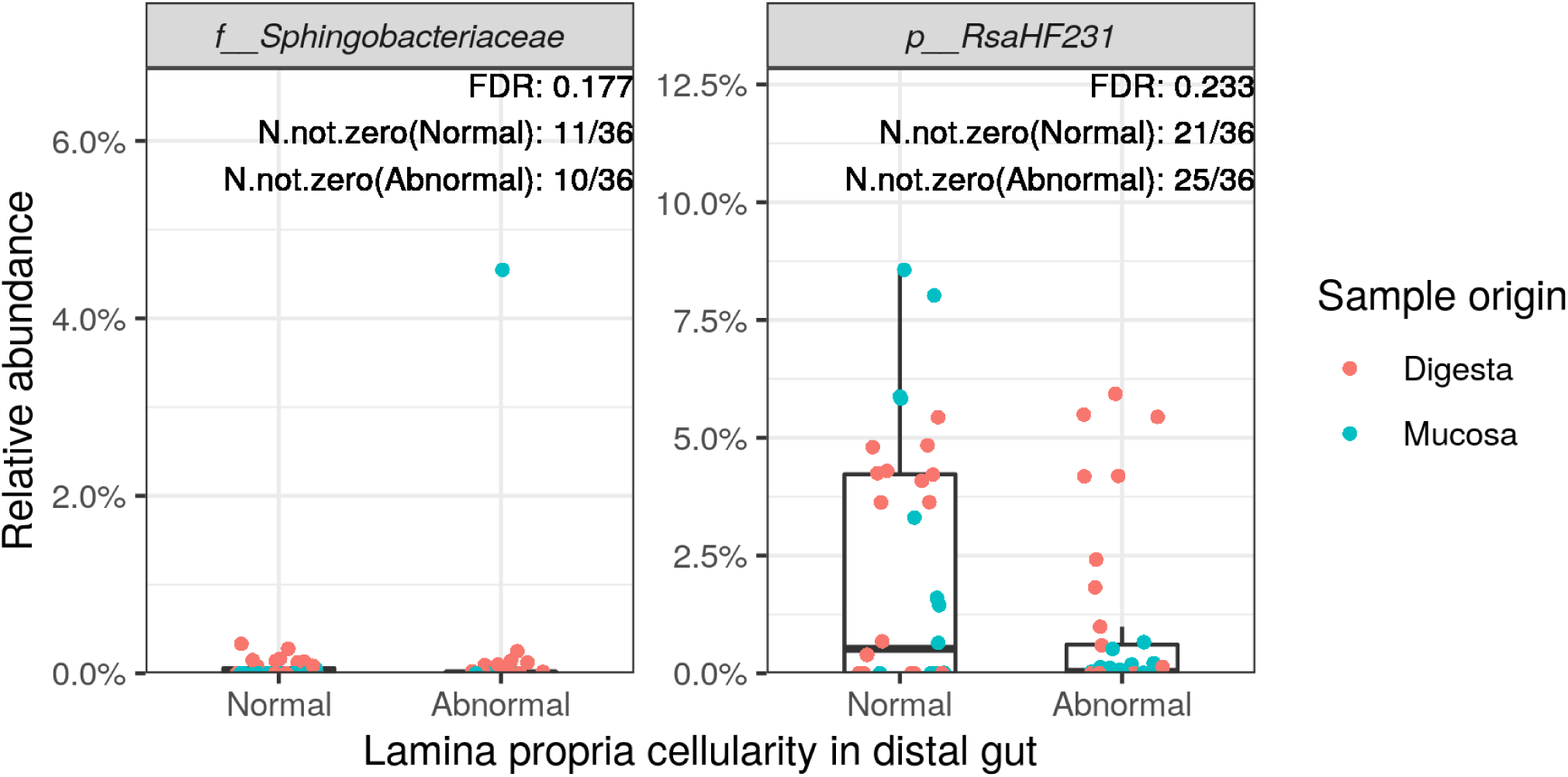
Microbial clades showing significant associations with histological scores on lamina propria cellularity in the distal intestine. *p*_, phylum; *f*_, family; FDR, false discovery rate; N.not.zero, number of non-zero observations.

**Fig. S6.**
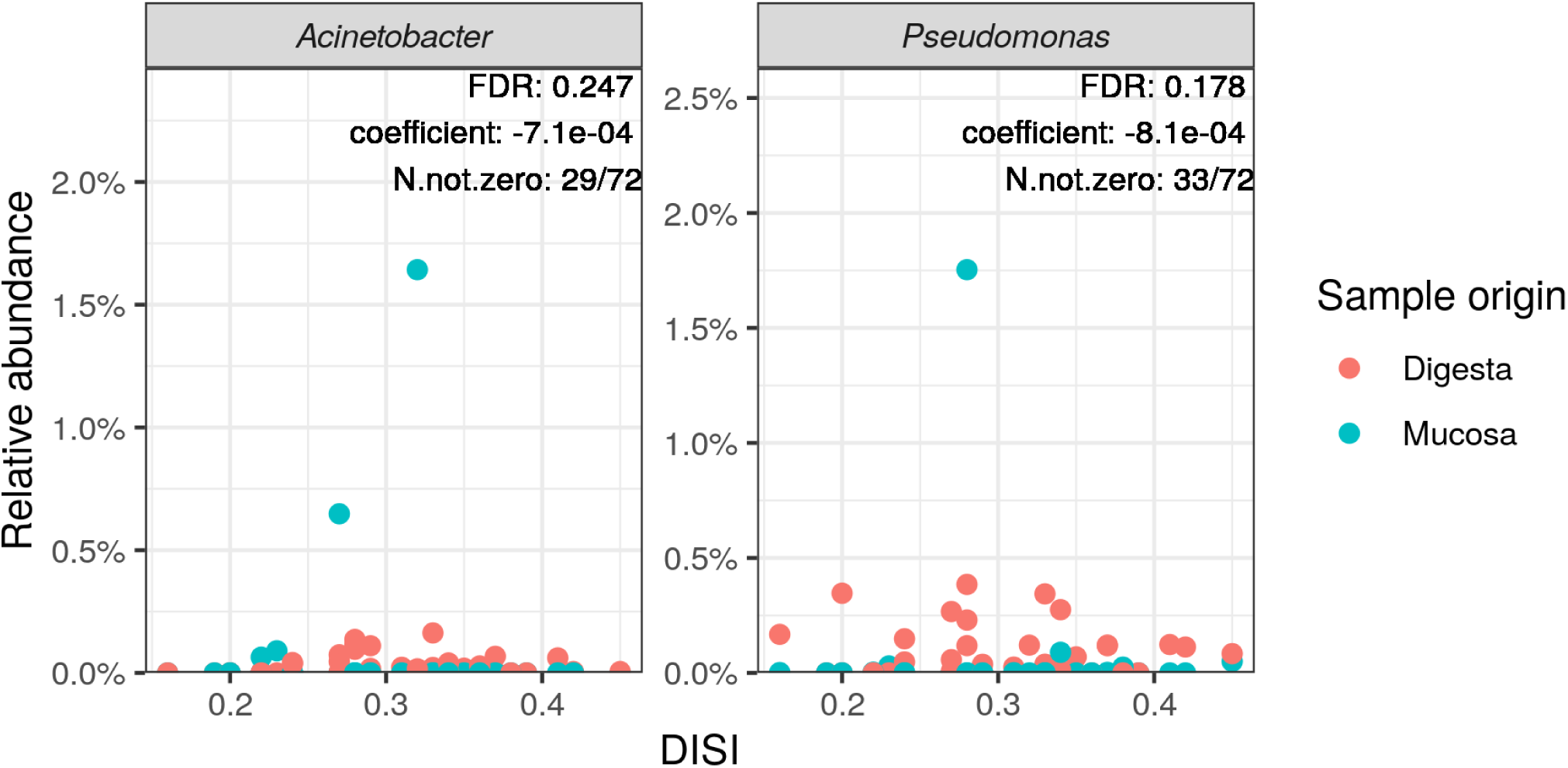
Microbial clades showing significant associations with distal intestine somatic index (DISI). FDR, false discovery rate; N.not.zero, number of non-zero observations.

**Fig. S7.**
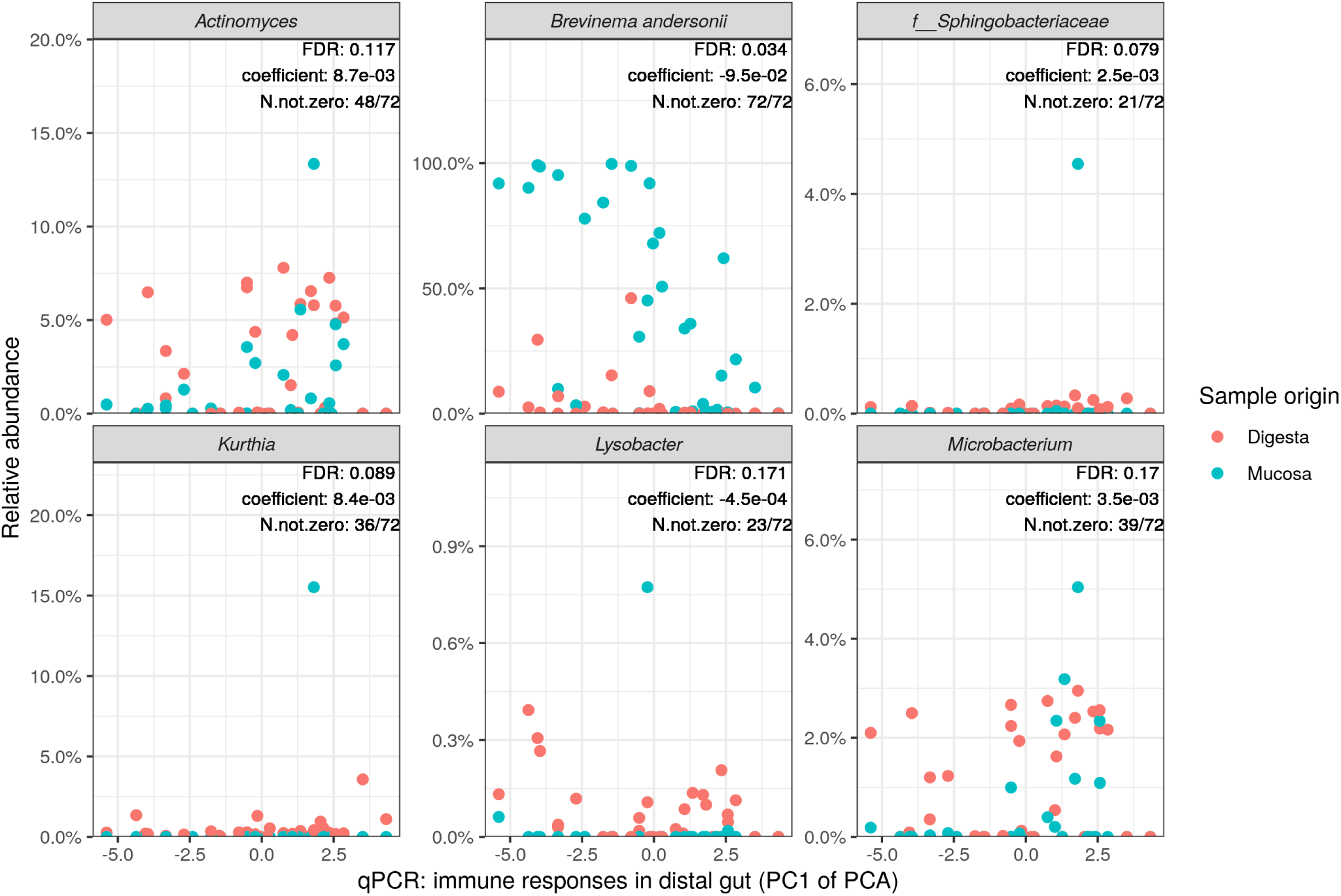
Microbial clades showing significant associations with immune gene expressions in the distal intestine. Since the expression levels of immune genes were highly correlated, we ran a principle component analysis (PCA) and used the first principle component (PC1) for the association testing to avoid multicollinearity and reduce the number of association testing. Note that the expression levels of immune genes decrease as the PC1 increases from left to right. Hence, a positive correlation coefficient denotes a negative association between the microbial clade and immune gene expressions, and vice versa. *f*_, family; FDR, false discovery rate; N.not.zero, number of non-zero observations.

**Fig. S8.**
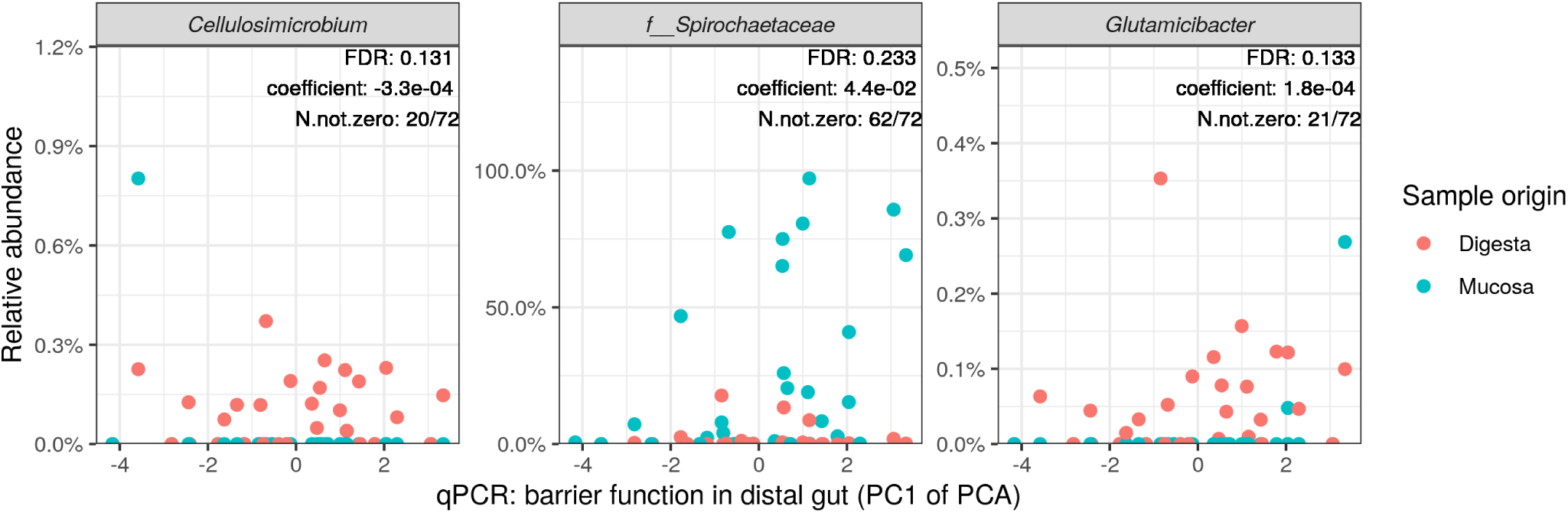
Microbial clades showing significant associations with expressions of barrier function related genes in the distal intestine. Since the expression levels of barrier function related genes were highly correlated, we ran a principle component analysis (PCA) and used the first principle component (PC1) for the association testing to avoid multicollinearity and reduce the number of association testing. Note that the expression levels of barrier function related genes decrease as the PC1 increases from left to right. Hence, a positive correlation coefficient denotes a negative association between the microbial clade and barrier function related gene expressions, and vice versa. *f*_, family; FDR, false discovery rate; N.not.zero, number of non-zero observations.

